# High-Fat Diet Induces Epigenetic and Metabolic Changes in Kisspeptin Neurons in Association with Obesity and Male Secondary Hypogonadism

**DOI:** 10.64898/2025.11.30.691258

**Authors:** Rui Fang, Rona S. Carroll, Ursula B. Kaiser

**Affiliations:** Division of Endocrinology, Diabetes and Hypertension, Brigham and Women’s Hospital and Harvard Medical School, Boston, MA, USA

**Author notes:** Communication and senior authors: Rui Fang, PhD,; Ursula B. Kaiser, MD.

**Keywords:** hypogonadism, obesity, diabetes mellitus, kisspeptin, 5hmC, TET2, AMPK, hyperglycemia, hydroxymethylation

## Abstract

**Background:** Obesity and type 2 diabetes mellitus (T2D) are major risk factors for male hypogonadism, a disorder with multi-system impacts on health. However, the mechanisms underlying obesity-associated male secondary hypogonadism remain poorly understood. Here, we aimed to dissect dysfunction of the hypothalamic–pituitary–testicular (HPT) axis and elucidate underlying epigenetic mechanisms, using a high-fat diet mouse model.

**Methods:** Male C57BL6 mice were fed standard chow or high-fat diet (HFD, 60% fat) for 16 weeks starting at age 6 weeks. Plasma testosterone levels, sperm counts, and gonadotropin responses to senktide and kisspeptin stimulation were assessed. HFD-induced transcription changes in the hypothalamic arcuate nucleus (ARC) were evaluted using bulk and single-cell RNA-sequencing. Genome-wide changes in 5-hydroxymethylcytosine (5hmC) were analyzed by hydroxymethyl-DNA immunoprecipitation sequencing (hMeDIP-seq). Functional relevance of 5hmC changes was evaluated by ectopic TET expression in an immortalized ARC Kiss1 neuron cell line and RT-qPCR.

**Results:** HFD-fed mice developed obesity, hyperglycemia, glucose intolerance, and insulin resistance, indicative of the development of a type 2 diabetes-like metabolic disorder. This was accompanied by low testosterone, reduced sperm counts, and unchanged basal luteinizing hormone (LH), consistent with obesity/T2D-associated male secondary hypogonadism, as observed clinically in humans. Impaired LH responses to senktide, a Kiss1 neuron activator, but not to kisspeptin itself, identified suppressed Kiss1 neuron function as a key mechanism underlying secondary hypogonadism. RNA-seq analysis revealed dysregulation of metabolic and neural pathways in Kiss1 neurons. hMeDIP-seq demonstrated widespread 5hmC alterations in the ARC of HFD-induced obese/diabetic mice, correlated with dysregulation of fatty acid metabolism, neuronal activity, and synapse function pathways. Ectopic TET expression *ex vivo* in Kiss1 neuronal cell lines restored 5hmC levels and upregulated key metabolic and neuronal genes that were repressed in the ARC of DIO mice

**Conclusion:** Our findings demonstrate that Kiss1 neurons are highly sensitive to diet and metabolic changes, and that obesity/diabetes-induced 5hmC modifications play a key role in dysregulating metabolic and neuronal pathways in Kiss1 neurons. These findings reveal a novel mechanism linking metabolic disturbances to reproductive dysfunction, through direct effects on Kiss1 neurons.

**Graphic Abstract:** 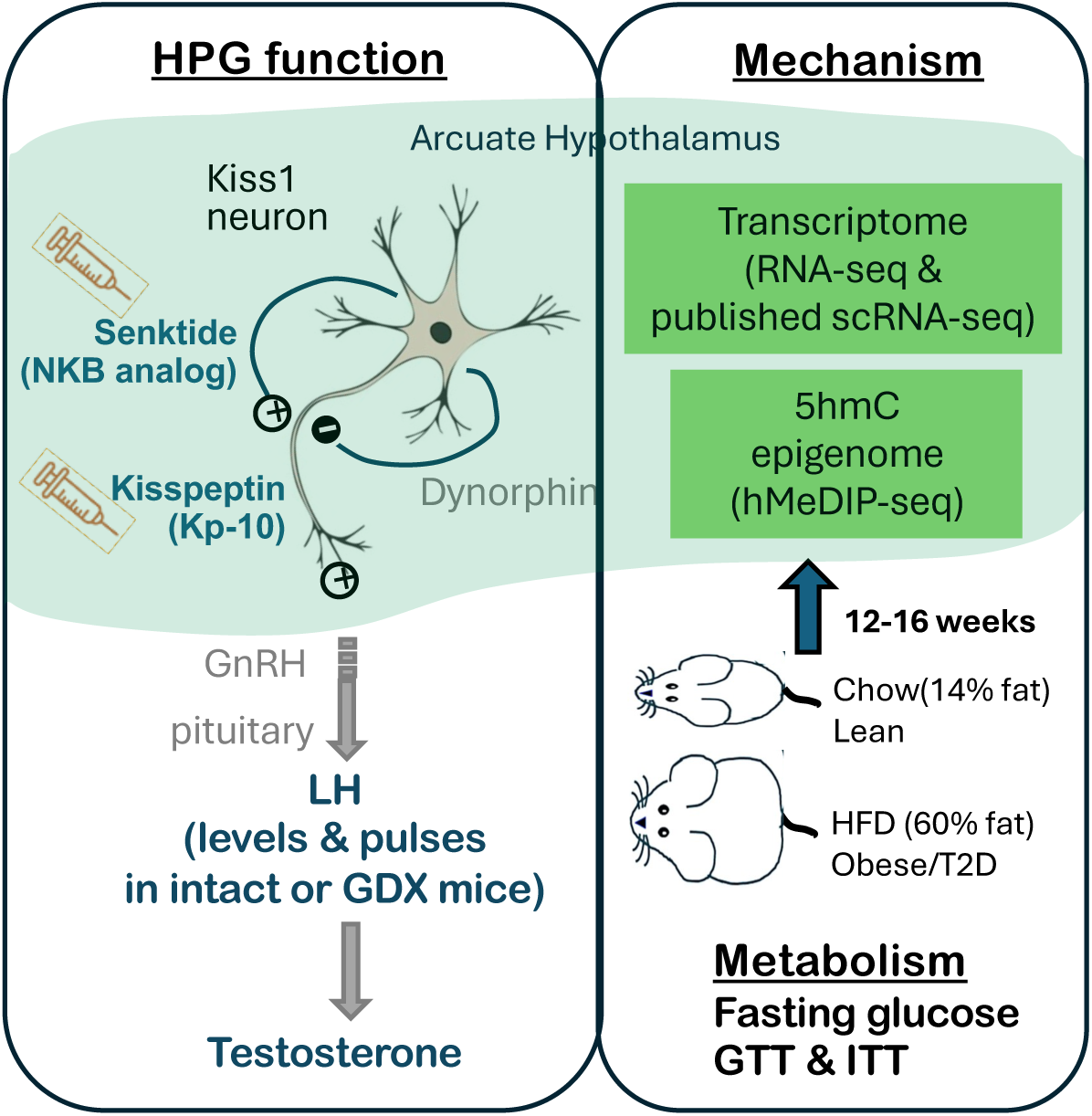

## Introduction

Obesity and type 2 diabetes mellitus (T2DM) are major risk factors for male hypogonadism, a disorder that affects 35–45% of obese non-diabetic adult males and 45–55% of obese diabetic adult males [1]. Male secondary hypogonadism, characterized by low testosterone (T) levels (total testosterone <8 nmol/L) with low or inappropriately normal luteinizing hormone (LH) levels (excluding that caused by structural hypothalamic–pituitary lesions) is commonly reversible through weight loss. However, significant increases in LH and testosterone, indicative of reactivation of the hypothalamic-pituitary-testicular (HPT) axis, typically requires substantial weight loss of 15% or more [2], which is challenging to achieve and sustain through lifestyle interventions alone. Low testosterone levels have multi-system impacts on health and are associated with fatigue, decreased libido, erectile dysfunction, depression, decreased bone density, loss of muscle mass, weight gain, increased risk of cardiovascular events, and increased mortality [3]. The rates of subfertility and infertility in young adult males are increasing [4] and are expected to worsen alongside the continued global rise of obesity and T2DM [5].

Kisspeptin (Kiss1) neurons are central to metabolic regulation of reproduction [6]. Kiss1 neurons in the hypothalamic arcuate nucleus (ARC) co-express neurokinin B (NKB), a Kiss1 neuron activator, and dynorphin, a Kiss1 neuron inhibitor—hence named KNDy neurons—forming an oscillator that governs the rhythmic activation of Kiss1 neurons and pulsatile release of kisspeptin, which subsequently activates hypothalamic GnRH neurons to, in turn, induce pulsatile LH (and FSH) secretion from pituitary gonadotrophs. The activity of Kiss1 neurons is regulated by upstream POMC and AgRP neuronal circuits, which exert opposing effects on food intake, energy expenditure, Kiss1 neuron activity, and reproductive function—with POMC neurons promoting and AgRP neurons inhibiting these processes. In addition, the energy sensors AMP-activated protein kinase (AMPK), mechanistic target of rapamycin (mTOR), and Sirtuin 1 (Sirt1) have been shown to play key roles in regulating Kiss1 neurons and pubertal timing in females on chow or high-fat diet [7, 8]. However, little is known about metabolite sensing in Kiss1 neurons in adults males with overnutrition.

5-hydroxymethylcytosine (5hmC), highly abundant in the brain, is increasingly recognized as a pivotal component of metabolism-mediated epigenome reprogramming that links environmental signals to adaptive changes in cellular function [9–12], including neurons [13, 14]. 5hmC is the oxidation product of 5-methylcytosine (5mC), mediated by the Ten Eleven Translocation family of DNA oxidases (TET1/2/3). During brain development, 5hmC plays a key role in maintenance and differentiation of neuronal progenitor cells [15]. In the adult brain, 5hmC and TET enzymes are important in neurogenesis, aging, and neurodegenerative diseases [16, 17]. Restoring 5hmC loss in the aged hippocampus through TET2 overexpression rejuvenates neurogenesis and enhances learning and memory [18]. We have previously identified the glucose-AMPK-TET2/5hmC signaling pathway, which links hyperglycemia to 5hmC loss and increased cancer risk associated with T2DM [11]. Therefore, we aimed to investigate the role of 5hmC in the epigenetic and metabolic regulation of Kiss1 neurons and HPT function.

We hypothesized that peripheral metabolites that cross the incomplete blood-brain barrier in hypothalamic arcuate nucleus induce epigenetic (5hmC), transcriptional, and metabolic changes in Kiss1 neurons, thereby regulating Kiss1 neuron function and hence gonadotropin secretion and reproduction. Through metabolic and neuroendocrine characterization, we demonstrate the development of male secondary hypogonadism in high-fat diet induced obese and diabetic mice, and show that Kiss1 neurons are sensitive to diet and metabolic changes and are key targets of the effects of metabolic disturbances. Importantly, we identified significant changes in the 5hmC methylome, which plays a key role in dysregulation of metabolism and gene transcription in Kiss1 neurons. These findings identify a novel molecular mechanism underlying crosstalk between Kiss1 neurons and metabolism, independent of upstream input from POMC and AgRP neurons.

## Methods

### Animal husbandry

Our study examined male mice exclusively, because obesity has significantly different pathophysiological effects on male and female reproductive health. Six-week-old C57BL/6J male mice were obtained from Jackson Laboratories. Animals were maintained at 22 ± 2°C on a 12-hour (h) light/dark cycle, with *ad libitum* access to water. Two experimental groups (n=8–12) were fed *ad libitum* standard chow (Purina 5053, 13.2% kcal from fat) or high-fat diet (Research Diets, D12492, 60% kcal from fat) for 3-4 months. Food consumption and body weight were monitored weekly. Mice were sacrificed using CO₂ and cervical dislocation without fasting. All animal studies were conducted as stated in the NIH *Guide for the Care and Use of Laboratory Animals* (National Academy Press, 1996) and approved by the IACUC of Brigham and Women’s Hospital and by the Harvard Medical Area Standing Committee on Animals in the Harvard Medical School Center for Animal Resources and Comparative Medicine.

### Mouse tissue collection

To collect the ARC, the brain was removed immediately after sacrificing the animal, and bilateral parasagittal cuts (1 mm lateral to the midline) were made to isolate the medial hypothalamus. Two coronal cuts were performed: one beneath the optic chiasm and another along the posterior edge of the median eminence. A horizontal cut, 1 mm deep, was then made to remove the targeted tissue, which was immediately frozen in liquid nitrogen.

Testes were isolated and weighed. To collect sperm, the epididymides were excised, minced, and the vas deferens were stripped of spermatozoa and cut into small pieces. The tissues were incubated in 2 ml of warm sperm collection medium (Dulbecco’s PBS with 14 mM DL-lactose, 16.7 µM pyruvate, 1 mM CaCl₂, and 0.5 mM MgCl₂) in a 35-mm petri dish at 37°C for 15–30 minutes to allow spermatozoa to swim out into the solution. The cell suspension was mixed gently, passed through a 40 µm cell strainer, and the petri dish and strainer were rinsed with 1 ml of sperm collection medium. The combined suspension was used for counting and calculation of the total number of sperm.

### Characterization of metabolism

To monitor the development of hyperglycemia, fasting glucose was measured for all experimental animals after a 4h fast and validated by a repeat test following a 14h overnight fast to ensure complete depletion of consumed food. Glucose levels from tail bleeds were measured using a glucometer (Agamatrix). For the insulin tolerance test, animals were fasted for 4h prior to an intraperitoneal (i.p.) injection of insulin (1.5 U/kg body weight; Humulin R U-100, Fisher Scientific). Glucose levels from tail bleeds were measured at 0 (pre-injection), 15, 30, 60, and 90 minutes post-injection. For the glucose tolerance test, animals were fasted for 14h overnight, and blood glucose levels were measured from tail blood at 0 (prior to injection), 15, 30, 60, 90, and 120 minutes (min) following intraperitoneal injection of glucose (1 g/kg in saline) using a glucometer (AgaMatrix). The average of two glucose readings at each time point was used for analysis. C57BL/6 mice are susceptible to development of glucose intolerance [19]. One mouse in the chow group consistently showed glucose intolerance and insulin resistance in GTT and ITT tests, and was excluded from the study. Obesity (as indicated by body weight) and hyperglycemia (determined by a 4h or 14h overnight fasting glucose) were confirmed in all HFD fed mice in the study.

### Gonadectomy

Bilateral gonadectomy was performed as described previously [20]. Briefly, a midline ventral incision (1 cm in length) was made under isoflurane inhalation anesthesia. The testes were removed bilaterally, and the vasculature was sutured to prevent internal bleeding. The body wall and skin were sutured to close the incision. Meloxicam analgesia was administered at the time of surgery and continued for three days postoperatively.

### Neuropeptide stimulation and LH secretion analysis

Animals were weighed prior to neuropeptide injection. Kp-10 peptide (Kisspeptin 110-119, Sigma-Aldrich K2644) was administered by intraperitoneal injection at a dose of 0.043 mg/kg, and Senktide (Sigma-Aldrich SML0764) was administered at 0.14 mg/kg. An interval of at least 7 days separated experiments. Tail blood (4 µL) collected at time 0 (prior to injection), and 15, 30, 45, and 60 minutes post-injection were diluted in 116 µl PBS-T (phosphate buffered saline, 0.1% Tween-20), immediately frozen on dry ice, and stored at -80°C. Plasma LH levels were quantified by ELISA as previously described [21]. Neuropeptide stimulation of LH secretion was calculated as Area Under Curve in arbitrary units using GraphPad Prism 7.

### LH pulses

Tail blood was taken every 8 minutes for 3h. Four µl tail blood were added 116 µl PBS-T (phosphate buffered saline, 0.1% Tween-20) and immediately frozen on dry ice. Plasma LH levels were quantified by ELISA in duplicates. LH peaks were identified by Matlab program as described previously [21]. LH amplitude was defined as the difference in LH levels between the peak and the nadir LH levels. Basal and peak LH levels were calculated as the average of the 3 lowest and 3 highest LH readings, respectively. LH production was calculated as the Area Under Curve using Prism 7.

### Plasma testosterone ELISA

Approximately 30 µl of tail blood was collected using a Microvette 100 Lithium Heparin Capillary Blood Collection Tube (Sarstedt) and centrifuged at 2,000 × *g* for 20 min at 4 °C. Plasma testosterone levels were measured in duplicate using the Mouse Testosterone ELISA Kit (Crystal Chem) according to the manufacturer’s instructions with minor modifications. Briefly, 5 µl of plasma or provided testosterone standards were mixed with 100 µl of Incubation Buffer and 50 µl of Enzyme Conjugate in each well, followed by shaking at 600 rpm at room temperature for 1 hour. After washing four times with 300 µl of Wash Buffer, 200 µl of Substrate was added and incubated for approximately 15 min at room temperature, protected from light. The reaction was stopped with 50 µl of Stop Solution before color saturation. OD450 was measured using a POLARstar Omega Microplate Reader (BMG Labtech) with OD630 as the reference wavelength. A standard curve (testosterone concentration vs. OD450) was generated using a semi-log fit, and plasma testosterone concentrations were calculated accordingly.

### RT-qPCR and RNA-seq

RNA and DNA were extracted from the same tissue sample using the Qiagen Allprep DNA/RNA micro kit per manufacture’s instructions. For RT-qPCR, first strand cDNA was synthesized from 0.5 µg RNA using iScript cDNA Synthesis Kits (Bio-rad). qPCR was performed using PowerUp SYBR Green Master Mix (Applied Biosystems). RT-qPCR primers were listed in **Supplemental Table 1**. RNA-seq library preparation and sequencing were conducted at Azenta Life Sciences (Burlington, MA) according to their standard protocol. Sequencing reads were filtered and trimmed using Trim Galore (https://github.com/FelixKrueger/TrimGalore). We aligned RNA-seq reads to the mm10 mouse genome using STAR 2.0 [22]. Counts per gene were generated by FeatureCounts using default parameters [23]. Differentially expressed genes were identified using EdgeR [24] and a cutoff of ≥1.5 fold change and p<0.05.

### 5hmC DNA immunoprecipitation sequencing

To reduce individual variability, biological replicates were generated by pooling equal amounts of DNA purified from the ARC of 3 mice. Two and 3 pools of biological replicates were generated for chow and HFD groups, respectively. hMeDIP-seq was performed as described previously [25]. Briefly, one µg of pooled DNA was sonicated into 100–200 bp fragments using a Covaris M2 sonicator and ligated with indexed adaptors using the NEBNext Ultra™ II DNA Library Prep Kit for Illumina (New England Biolabs) according to the manufacturer’s instructions. Barcoded DNA (50 µl in TE buffer: 10 mM Tris pH 8.0, 1 mM EDTA) was denatured at 98°C for 10 minutes, cooled rapidly in an ice water bath for 10 minutes and pooled in equal amounts (quantified by qPCR) across samples. Two µg of pooled, barcoded DNA and 4 µg of anti-5hmC antibody (Active Motif 39791) in 300 µl of 1× IP buffer (20 mM phosphate buffer pH 7.0, 0.14 M NaCl, 0.1% Triton X-100) were incubated overnight at 4°C. The DNA-antibody complex was immunoprecipitated by incubating with 30 µl of Dynabeads Protein A beads (Invitrogen) at 4°C for one hour and eluting in 1% SDS after four washes with 1× IP buffer. Enriched DNA was purified using the Qiagen MiniElute PCR Purification Kit and amplified with 12 cycles of PCR using the KAPA HiFi Library Amplification Kit (Roche). Illumina sequencing generated a minimum of 35 million paired end 75 bp reads per sample. hMeDIP-seq reads were aligned to the mm10 mouse genome using Tophat2.0 [26] after being filtered and trimmed using Trim Galore. hMeDIP-seq profiles were generated using Deeptools bamCoverage [27], normalized by total reads and visualized using the UCSC genome browser. Regions with significant 5hmC changes were identified using PePr [28] using default parameters and filtered with a cutoff of ≥1.5 fold change and p<0.001 (all DMRs identified had FDR<0.05). DhMR-associated genes were identified using ChIPseeker [29]. Gene Ontology analysis and KEGG pathway enrichment analysis were performed using DAVID [30] and GSEA [31].

### Statistics

Values are presented as mean ± SEM. Student’s *t*-test was used when comparing two groups. One-way ANOVA followed by Tukey’s HSD test was used for multi-group analyses. p<0.05 was considered significant. Box whisker plots were generated by R. The line inside the boxplots denotes the median, the box represents the interquartile range (IQR), and the whiskers extend to the minimum and maximum values within 1.5 times the IQR from the lower and upper quartiles. Outliers beyond this range are displayed as individual points.

## Results

### High-fat diet induces obesity, diabetes, and secondary hypogonadism

We fed C57BL/6 male mice a high-fat diet (HFD, Rodent Diet D12492, 60% fat, n=12) or standard chow (chow, Purina 5053, 13.2% fat, n=7), beginning at age 6 weeks. We monitored fasting glucose, glucose tolerance, and insulin sensitivity of both groups, and characterized HPT axis physiology following significant weight gain and the development of hyperglycemia and insulin insensitivity in the HFD group. We collected ARC tissue for RNA sequencing (RNA-seq) and hydroxymethylated DNA immunoprecipitation coupled with deep sequencing (hMeDIP-seq) analysis after 16 weeks on HFD.

The body weight of mice on HFD was significantly higher than that of chow-fed mice as early as one week after starting the diet (26.3±0.5g vs 22.8±0.2g in HFD and chow fed mice, respectively; mean±standard error of the mean [SEM]; p<0.001), 56% heavier after 10 weeks on HFD (46.1±1.0 g vs 29.5±0.4 g, p<10^-9^), and 65% heavier at the time of sacrifice after 16 weeks on HFD (48.3±0.8 g vs 29.3±0.6 g, p<10^-10^) (**Figure 1A**). Elevated fasting blood glucose levels (after a 4h fast) in high fat diet-induced obese (DIO) mice were observed as soon as 4 weeks on HFD (261.8±11.1 vs 226.0±9.5 mg/dL in HFD and chow fed mice, respectively; 16% increase; p<0.05), coinciding with a 37% heavier body weight at this time point (35.1±0.9 g vs 25.6±0.3 g body weight after 4 weeks of HFD or chow, respectively; p<10^-6^). After 10 weeks on HFD, 4-hour fasting glucose was 30% higher in DIO mice (282.9±15.0 mg/dL vs 218.0±12.4 mg/dL in HFD and chow fed mice, respectively; p<0.01). Glucose levels after a 14h fast were 49% higher (186.8±4.3 mg/dL vs 125.1±4.0 mg/dL, p<10^-7^, **Figure 1B**). DIO mice had higher blood glucose levels in glucose tolerance tests (**Figure 1C**, p<0.05). Glucose clearance was significantly delayed and remained high in insulin tolerance tests in DIO mice (p<0.001; **Figure 1D**), indicating insulin resistance.

**Figure 1.**
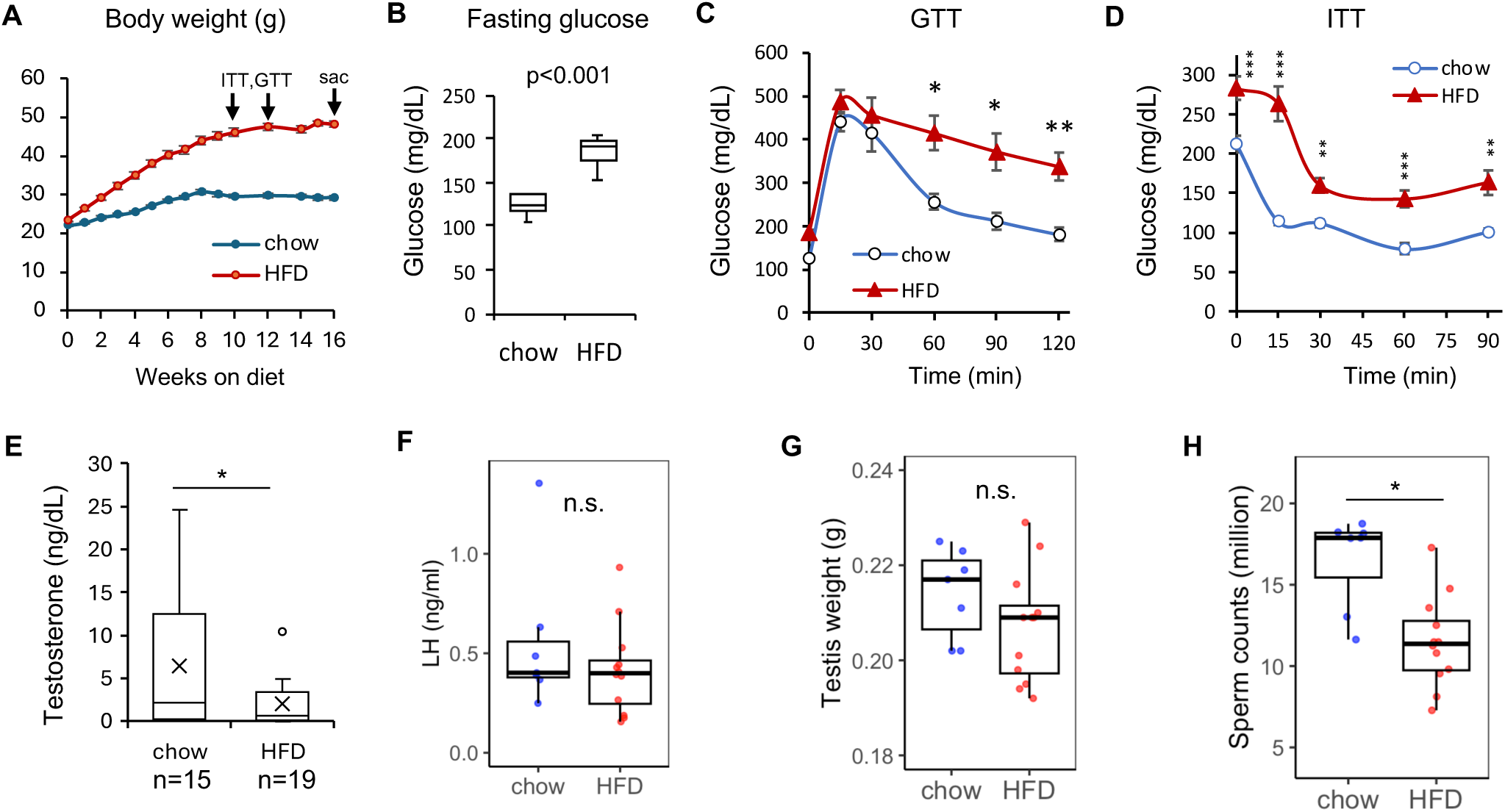
High-fat diet (HFD) fed mice as a model of male obesity-associated secondary hypogonadism. **A.** Body weight of male mice on standard chow (n=7) or high-fat diet (HFD, 60% fat/cal, n=12), starting at age 6 weeks. Mice on HFD were significantly heavier beginning after 1 week on HFD (Student’s *t*-test, p<0.001). Mice had insulin tolerance test (ITT) at age 10 weeks, glucose tolerance test (GTT) at age 12 weeks, and were sacrificed (sac) at age 16 weeks, with HFD continued until time of sacrifice (sac). Error bars, SEM. **B.** Boxplot of Blood glucose levels in chow (n=7) and HFD (n=12) fed mice after a 14h fast. **C.** Glucose tolerance test (GTT; 1 g/kg glucose ip after overnight fast) in chow (n=7) and HFD (n=12) fed male mice. Error bars, SEM. **D.** Insulin tolerance test (ITT; 1.5 U/kg insulin ip after 4h fast) in chow (n=7) and HFD (n=12) fed male mice. Error bars, SEM. **E.** Boxplot of plasma testosterone (T) levels in HFD (n=19) and chow (n=15) fed mice. Open circle denotes outlier beyond the whisker mark. **F.** Baseline plasma LH in chow (n=7) and HFD (n=12) mice. **G, H.** Boxplot of testis weight (**G**) and sperm counts (**H**) of mice on chow (n=7) or HFD (n=12). P values in figures, Student’s *t*-test; *, p<0.05; **, p<0.01; ***, p<0.001; n.s., not significant. In boxplots in **E**-**H**, the box boundaries represent the 25th and 75th percentiles; line within the box indicates the median; whiskers extend to 1.5 × interquartile range (IQR) beyond the box. Open circle in **E** represent outliner outside of 1.5 × the interquantile range. All data points are shown in **F-H**

We next measured testosterone and LH levels to evaluate HPT function. Testosterone levels were significantly lower in DIO mice (p<0.05; **Figure 1E**). In healthy mice, low testosterone is expected to activate the HPT axis, resulting in elevated plasma LH levels. However, LH levels in DIO mice were not different from the chow group (**Figure 1F**). There was no change in testis weight (**Figure 1G**). Sperm counts were significantly reduced in DIO mice (p<0.05; **Figure 1H**). Low T levels accompanied by an absence of increase in LH in diet-induced obese/diabetic mice mirrors clinical findings in patients with obesity and type 2 diabetes mellitus, indicating secondary (i.e., central) hypogonadism.

### Dysregulation of LH production in gonadectomized (GDX) DIO mice

Different T levels in the chow and HFD groups may affect central HPT axis activity through altered negative feedback. The higher T levels in the chow control group are expected to exert a greater negative feedback effect on the central HPT axis, compared to the effects of the lower T levels in the DIO mice, potentially obscuring inhibited HPT activity in DIO mice. Suboptimal testicular function associated with obesity may be attributed to direct effects of factors such as oxidative stress and chronic inflammation, in addition to potential indirect effects from a suppressed central HPT axis[32]. To overcome the effects of differences in testicular function and accurately assess central HP function, we performed gonadectomy (GDX) in a cohort of male mice after 12 weeks of HFD, and continued the same diet after surgery. Prior to surgery, the HFD fed mice were obese (46.7±1.1g vs 29.2±0.75g, 61% heavier, p<10^-5^) and had higher fasting glucose levels (overnight fasting glucose 194.4±8.0mg/dL vs 138.7±4.5mg/dL, p<0.001, 40% increase). A significant increase in LH levels was observed 24 hours post-GDX (day 1) in chow-fed lean mice, as expected (**Figure 2A)**. The increase in LH was significantly delayed in castrated HFD-fed and chow control mice (**Figure 2A**, ANOVA followed by Tukey’s HSD test, p<0.01). There were no significant differences between HFD-fed and chow control mice at day 0 (prior to surgery) or from day 2 to day 27 post-surgery (p>0.05; **Figure 2A**).

**Figure 2.**
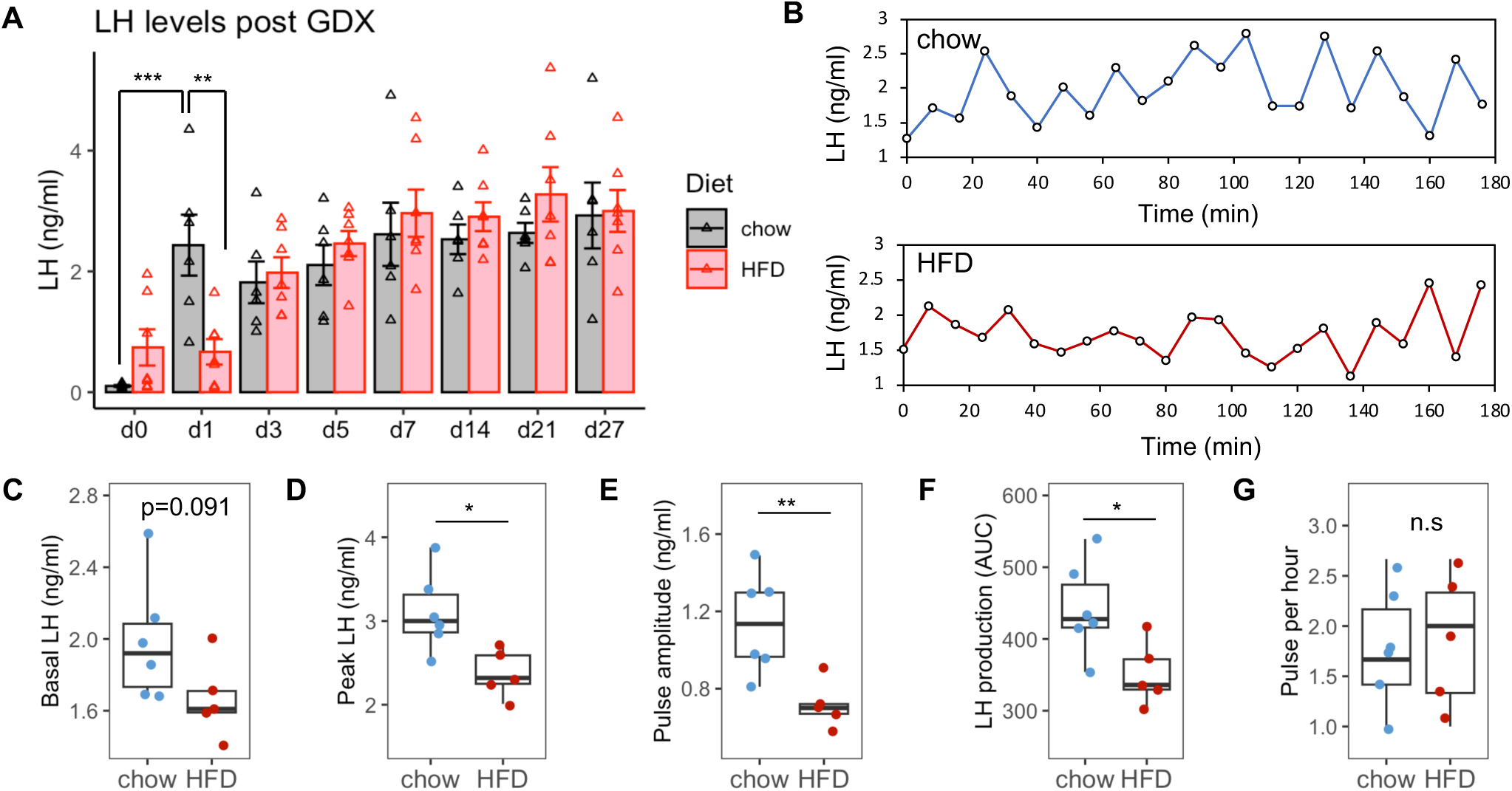
LH pulses in castrated male mice fed high-fat diet or standard chow. **A.** The LH increase after gonadectomy (GDX) was delayed in HFD mice (n=7) compared chow control group (n=6). ANOVA, p<0.05. **, Tukey’s HSD test, p<0.01; ***, Tukey’s HSD test, p<0.001. Error bars, SEM. **B.** Representative LH pulses of GDX chow and HFD mice. **C.** Basal LH levels showed a trend to be lower in HFD GDX mice, but did not reach statistical signficiance (p>0.05). Chow, n=6; HFD, n=5. **D-F.** Mean peak LH value (D), LH pulse amplitude (E) and total LH production (F; AUC) were significantly lower in HFD GDX mice than in chow GDX mice. *, p<0.05; **, p<0.01; chow, n=6; HFD, n=5. **G.** There was no significant difference in LH pulse frequency between GDX male HFD and chow groups (n.s., not significant; by Student’s *t*-test); chow, n=6; HFD, n=5. In boxplots in **C-G**, the box boundaries represent the 25th and 75th percentiles; line within the box indicates the median; whiskers extend to 1.5 × IQR beyond the box; all data points are shown.

To further characterize central HPT function, we assessed LH pulsatility in chow and DIO GDX mice by measuring plasma LH levels in tail bleeds every 8 minutes for 3h on days 41-44 after GDX. The mean body weight of HFD and chow groups were 47.2±4.0 g (n=5) and 27.3±1.2 g (n=6), respectively, which did not differ significantly from pre-operative body weights. We observed “flatter” LH curves (i.e., less variability in LH levels) in DIO mice compared to controls (representative LH frequent sampling data shown in **Figure 2B**). While we did not observe significant differences in the mean nadir baseline LH levels (**Figure 2C)**, the mean peak LH levels were significantly lower in DIO mice compared to controls fed normal chow (p<0.05, **Figure 2D**). The mean LH pulse amplitude, defined as the difference between peak and nadir LH levels, was also significantly lower in DIO mice (p<0.01, **Figure 2E**). Total LH production, calculated as area under the curve (in arbitrary units), was also significantly lower in DIO mice (p<0.05, **Figure 2F**). There was no difference in LH pulse frequency between the two groups (**Figure 2G**).

Significant weight loss during the first week post-GDX was observed in the HFD group (mean 9.0 g/19.0% bodyweight loss, p<0.001), which was more pronounced than in the chow group (mean 2.1 g/8.2% body weight loss, p=n.s.) (**Figure 3A**). Weight loss in the HFD group was accompanied by normalization of blood glucose levels as early as 5 days post-GDX (**Supplemental Figure S1**). On day 27, when no differences in LH levels were detected between the intact chow and HFD groups (**Figure 2A**), HFD GDX mice had elevated blood glucose levels compared to chow-fed mice (209.3±11.9 mg/dL vs 187.7±5.1 mg/dL, p<0.01 by Student’s *t*-test; **Supplemental Figure S1**), and a mean body weight of 40.4±2.4 g which was significantly lower than the pre-surgery (day 0) body weight (14.4% lower, p<0.05; **Figure 3A**). On day 72 post-GDX, when body weight and glucose levels had returned to pre-GDX levels (**Figure 3A**, mean body weight 47.3±2.5 g and 29.3±0.7 g in HFD and chow groups, respectively; **Figure 3B**, mean 4h fasting glucose 233.1±7.8 mg/dL vs 174.3±2.7 mg/dL for HFD and chow groups, respectively; p<0.05), we observed a significant decrease in plasma LH levels in long-term GDX DIO mice compared to chow mice (**Figure 3C**, 1.32±0.09 vs 2.36±0.12 ng/ml, p<0.001).

**Figure 3.**
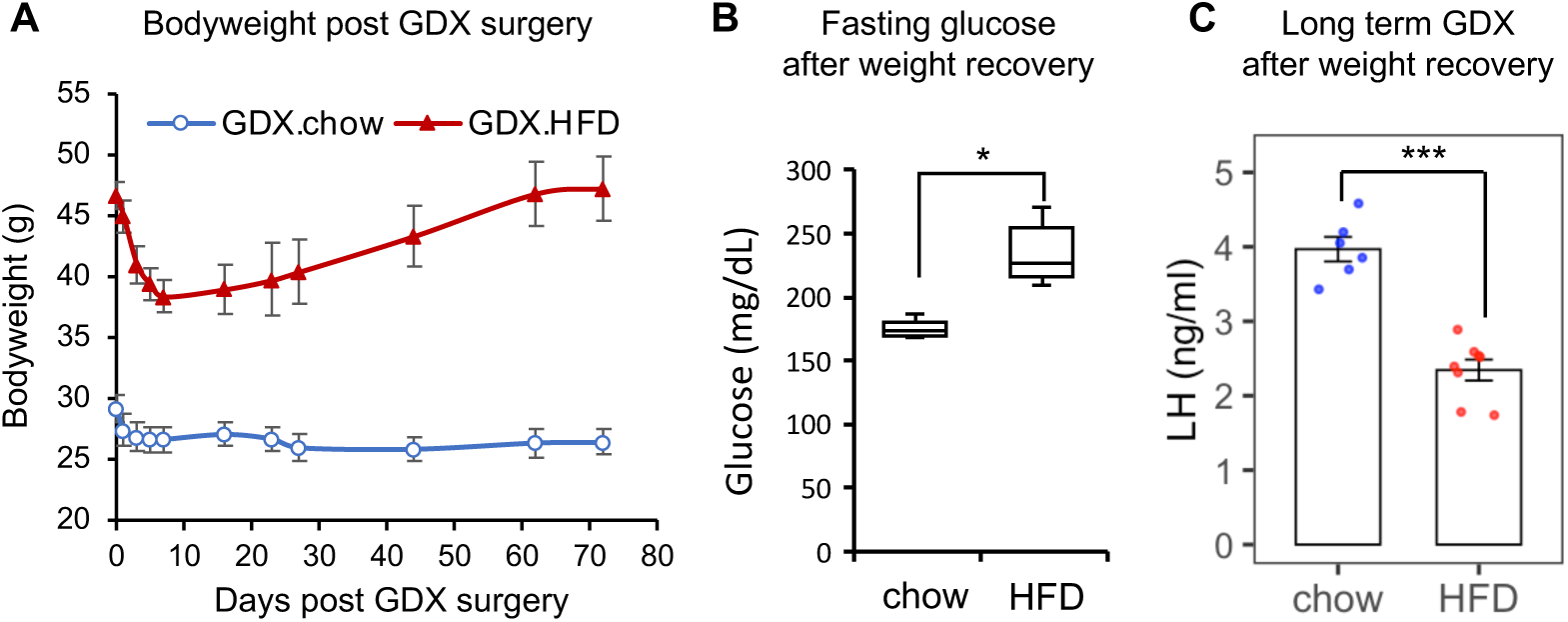
Plasma LH levels are lower in long-term GDX male obese diabetic mice. **A.** Bodyweight loss and recovery after GDX in mice fed HFD or standard chow diet. Error bars, SEM. **B.** Fasting plasma glucose levels after a 4h fast, on day 72 post-GDX. Chow, n=6; HFD, n=8. *, p<0.05. The box boundaries in boxplot represent the 25th and 75th percentiles; line within the box indicates the median; whiskers extend to 1.5 × IQR beyond the box. **C.** Plasma LH levels in long-term GDX (day 72 post GDX surgery) mice fed chow or HFD. The mean of two plasma LH measurements taken 8 minutes apart were plotted. Error bars, SEM. Chow, n=6; HFD, n=8. ***, p<0.001.

Thus, after GDX, the delayed increase in plasma LH, reduced LH pulse amplitude, and lower LH levels in GDX HFD fed mice compared to mice fed standard chow indicate suppressed central HPT function in obese diabetic mice.

### LH secretion in response to senktide, but not kisspeptin, is reduced in intact DIO mice compared to controls

While suppression of HPT function by overnutrition is consistently observed in clinical studies in men and in HFD animal models, the reported results from efforts to pinpoint the level of the HPT dysfunction have been inconsistent. Some studies indicate GnRH neuron dysfunction [33], while others report Kiss1 neuron dysregulation with preserved GnRH neuron function [34–37]. To gain additional insight into the locus of HPT dysfunction in our HFD-fed GDX mouse model, we performed kisspeptin and senktide stimulation tests (**Figure 4A**). We observed similar levels of basal LH and peak stimulated LH after a single intraperitoneal injection of kisspeptin-10 (kp-10) in intact lean and DIO mice (0.043 mg/kg, **Figure 4B**), suggesting normal GnRH neuron and pituitary gonadotrope function and responsiveness to kisspeptin. Intraperitoneal injection of senktide, a neurokinin B analog and Kiss1 neuron activator, induced a robust LH surge, peaking at 15 minutes post-injection. LH levels 30 and 60 minutes after senktide administration were significantly lower in DIO mice compared to chow mice (**Figure 4C**). The total LH production in response to senktide stimulation (AUC) was also significantly lower in DIO mice (**Figure 4C**, p<0.05), indicating reduced Kiss1 neuron activation in response to a neurokinin B agonist.

**Figure 4.**
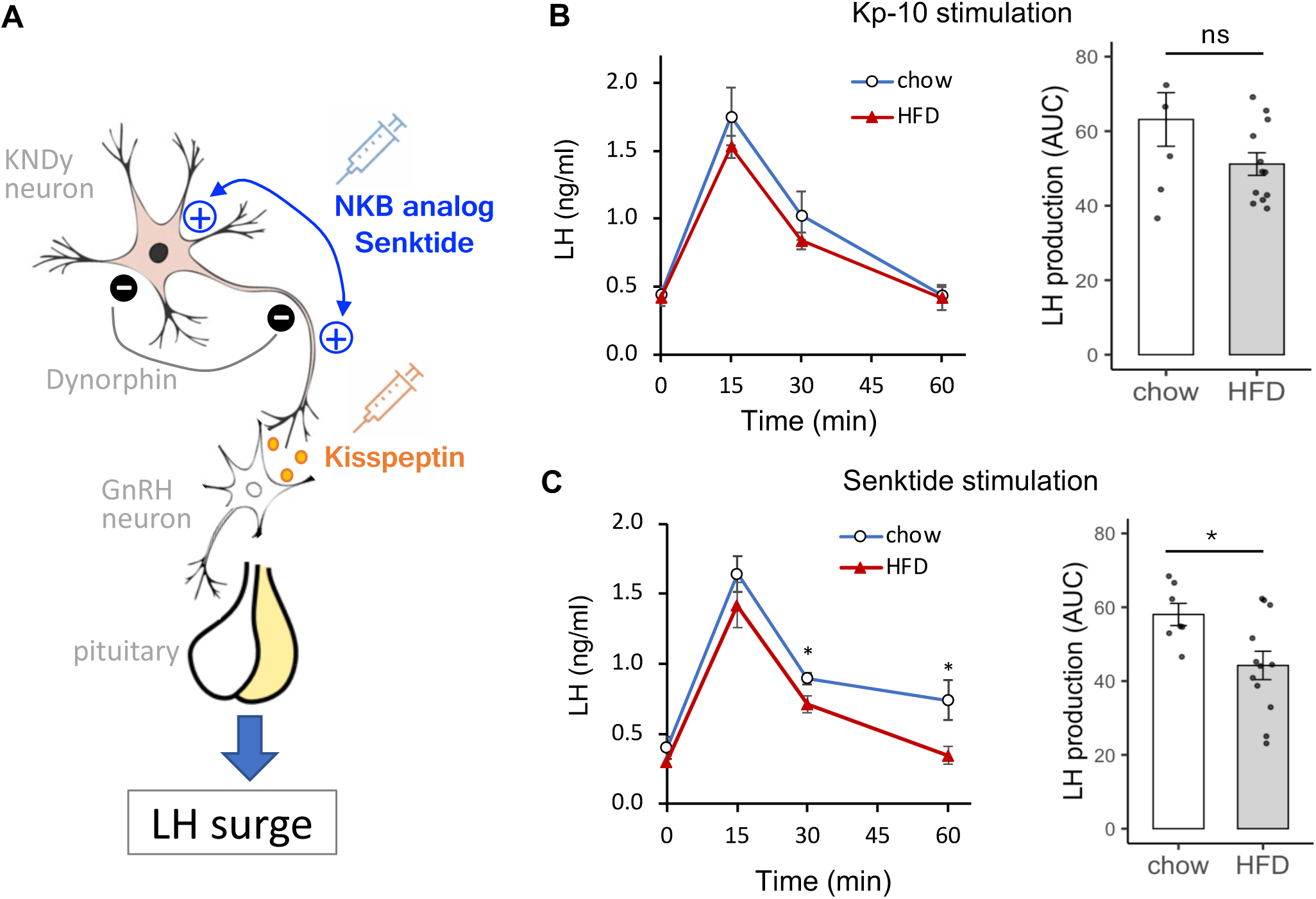
LH secretion in response to senktide and kisspeptin in intact males fed HFD or chow. **A.** Schematic of neuropeptide stimulation to dissect central HPT axis function. **B.** Kisspeptin (0.043 mg/kg ip)-stimulated increase in plasma LH (left) and relative cumulative LH production (AUC, right) in intact chow (n=6) and HFD (n=12) fed male mice. ns, not significant; Error bars, SEM. **C.** Senktide (0.14 mg/kg ip)-stimulated increase in plasma LH (left) and relative cumulative LH production (AUC, right) in intact chow (n=6) and HFD (n=9) male mice. *, p<0.05. Error bars, SEM.

### Metabolic and neuronal gene networks are reprogrammed in Kiss1 neurons in the ARC in DIO mice

To elucidate the mechanisms underlying Kiss1 neuron dysfunction in obese diabetic mice, we performed RNA-seq transcriptome analysis of the ARC from chow (n=3) and DIO (n=4) mice. The transcriptomes of the two groups demonstrated significant differences, showing clear separation in the principal component analysis plot (**Figure 5A**). We identified 196 up-regulated genes and 480 down-regulated genes, using a cut-off of ≥1.5-fold change and p<0.05. Many key fatty acid and lipid metabolism related genes were down-regulated in DIO ARC, such as fatty acid binding proteins *Fabp1* and *Fabp7*, apolipoproteins *Apoa1*, *Apoa5* and *Apoh*, and orphan nuclear receptor 4a1 (*Nr4a1*) (blue dots in **Figure 5B**). Genes encoding glucose transporter type 1 (*Slc2a1*), acetylcholine transporter (*Slc18a3*), high affinity choline transporter (*Slc5a7*), choline acetyltransferase (*Chat*), and the Rho2 subunit of the GABA type A/C receptor complex (*Gabrr2*) were upregulated (green and red dots in **Figure 5B**). No significant changes in *Kiss1*, *Kiss1r*, *Tac2* or *Pdyn* gene expression were observed in the ARC by RNA-seq or RT-qPCR analysis of intact (**Supplemental Figure 2**) or GDX mice (data not shown).

**Figure 5.**
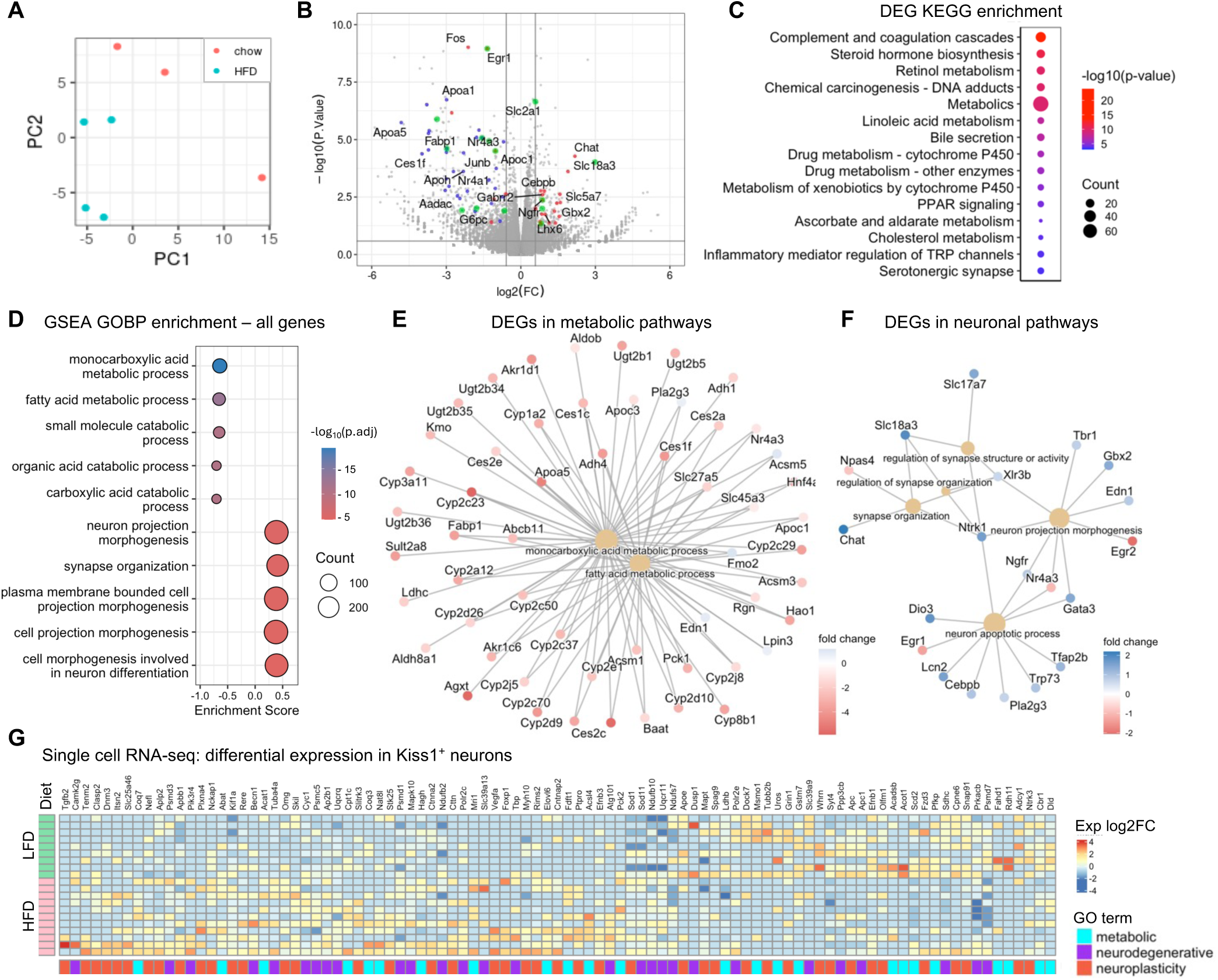
Dysregulation of metabolic and neuronal gene pathways in the ARC of HFD mice. **A.** Principal component plot of gene expression in the ARC of mice fed either chow or HFD. Red, three biological replicates of chow ARC; blue, four biological replicates of HFD ARC. **B.** Volcano plot of gene expression changes. Blue, relates to fatty acid metabolism; red, neuron function; green, glucose metabolism. Horizontal line, p=0.05; vertical lines, |log_2_FC|=0.585. **C.** Top fifteen enriched KEGG pathways of DEGs in HFD ARC compared to chow diet ARC. **D.** GSEA analysis of gene expression changes in HFD and chow ARC. The top five suppressed and activated GOBP pathways are shown. **E-F.** DEGs in top repressed metabolic pathways (**E**) and top activated neuronal GOBP pathways (**F**). Red and blue reflect down- and up-regulation, respectively. Center yellow nodes denote associated GOBP pathways. **G.** Heatmap of unsupervised clustering of differential gene expression in ARC Kiss1 neurons revealed by single-cell RNAseq after 1 week HFD and LFD treatment[40]. Each row represents a Kiss1-positive neuron (≥1 reads mapped to *Kiss1* gene). A relaxed cutoff (|FC|>=1.5 & p<0.2) were used to select differentially expressed genes due to scarcity of Kiss1^+^ neurons and low reads coverage per cell of single-cell RNAseq data.

KEGG pathway analysis of differentially expressed genes (DEGs) revealed enrichment of metabolism-related pathways, including the general KEGG metabolic pathways, linoleic acid metabolism, PPAR signaling, and cholesterol metabolism, as well as the inflammatory mediator regulation of TRP channels and serotonergic synapse pathways (top 15 pathways affected shown in **Figure 5C**). Using Gene Set Enrichment Analysis (GSEA), which uses whole-genome changes in expression to identify coordinated changes of functionally related genes [31], we observed that metabolism pathways were the most significantly suppressed and the neuronal pathways were the most upregulated Gene Ontology Biological Process (GOBP) in DIO mice (top 5 shown in **Figure 5D**). Monocarboxylic acid metabolic process is a key energy source for the brain that involves the metabolism of pyruvate, lactate, acetate, and short-chain fatty acids. It has considerable overlap with fatty acid metabolic process. The majority of the DEGs (≥1.5-fold change and p<0.05) related to these two processes were down-regulated in DIO mice (**Figure 5E**, left panel).

The top upregulated GOBP neuronal pathways include the Neuron Projection Morphogenesis Pathway and the Synapse Organization Pathway, which are essential for neuronal plasticity in the adult brain, with the former involved in axonal and dendritic projection and the latter in the formation and maintenance of synapses (top 5 shown in **Figure 5B**; reduce representation in **Supplemental Figure 4**). Most DEGs associated with these pathways were up-regulated (**Figure 5E**). These observations suggest that HFD-induced obesity and type 2 diabetes mellitus have significant impact on the transcription of neuronal and metabolic gene networks, suggesting significant changes in neuron metabolism and function in association with increased activities in neuroplasticity.

The ∼3000 Kiss1 neurons in the ARC account for a small fraction of the cells in the hypothalamic tissue we collected for bulk RNA sequencing. To gain insight into Kiss1 neuron-specific effects, we reviewed published single-cell and single-nucleus RNA-seq datasets[38–40]. We found that sequencing reads mapped to the *Kiss1* gene were too sparse in single nucleus RNA-seq datasets to identify Kiss1^+^ neurons based on *Kiss1* gene expression alone[38, 39]. The Campbell study performed single cell Drop-seq analysis on micro-dissected ARC-ME in male mice fed a low-fat diet (Ch10, 10% fat) or high-fat diet (HFD, 60% fat) for one week [40]. In this dataset, we identified 9 Kiss1 neurons in the groups of low-fat diet fed mice, and 11 Kiss1 neurons in high-fat diet fed mice, respectively, with ≥1 read counts mapped to the *Kiss1* gene. The sparse numbers of reads and low numbers of cells limited statistical power to identify differentially expressed genes. Using a relaxed cutoff of |FC|≥1.5 and p<0.2, we identified 340 up-regulated and 383 down-regulated genes in HFD Kiss1 neurons, compared to LFD. Strikingly, the top enriched KEGG gene pathways of DEGs in Kiss1 neurons are predominantly associated with neuronal function and metabolism (**Supplemental Figure 5**). Nine of 15 identified KEGG gene pathways are related to neuronal function, including Amphetamine Addiction, Long-Term Potentiation, and all pathways within the neurodegenerative disease subcategory—namely Alzheimer’s Disease, Parkinson’s Disease, Amyotrophic Lateral Sclerosis, Huntington’s Disease, Spinocerebellar Ataxia, and Prion Disease (**Supplemental Figure 5**). DEGs related to neuronal plasticity and neurodegenerative diseases were predominantly upregulated in HFD Kiss1 neurons, with twice as many genes up-regulated as down-regulated (**Figure 5G**). Five of the 15 top enriched KEGG pathways are related to metabolism, including pyruvate metabolism, insulin secretion, metabolic pathways, non-alcoholic fatty liver disease, and carbon metabolism (**Supplemental Figure 5**). There are ∼110 out of the 723 DEGs related to metabolic pathways. A subset related to lipid metabolism are shown in **Figure 5G** (cyan). A similar analysis of GnRH neurons in the Campbell dataset identified a small number of differentially expressed metabolic genes (data not shown)[40]. These observations indicate that Kiss1 neurons in the ARC are likely to be susceptible to effects of HFD.

Taken together, single-cell and bulk RNA-seq studies indicate that HFD markedly impacts the metabolism and function of Kiss1 neurons. Notably, only a small number of genes are consistently dysregulated in Kiss1 neurons from short-term HFD-treated mice[40] and in the ARC of long-term HFD-fed mice in our studies. This is not unexpected, given the significant physiological differences between the groups: mice with short-term HFD feeding exhibit grossly normal metabolism, whereas long-term HFD-fed mice (DIO model) have substantial metabolic dysfunction, including obesity, hyperglycemia, and insulin resistance. Interestingly, immediate early response genes, including *Dusp1*, *Fos*, *Fosb*, *Egr1*, and *Nr4a1*, were consistently downregulated in the HFD group across both single-cell and bulk RNA-seq datasets. These genes are known to play key roles in synaptic plasticity and neuronal activity [41].

### HFD induces 5hmC changes in the ARC in key metabolic and neuronal genes

We found that expression of *TET1* and *TET2* in the human hypothalamus correlated strongly with *KISS1*, *TAC3* (encoding neurokinin B in humans), and *TACR3* (encoding the neurokinin B receptor) gene expression in the Genotype-Tissue Expression (GTEx) database. *TET3* expression correlates positively with *TAC3* and *TACR3* but not *KISS1* expression (**Supplemental Figure 7**). Single-cell RNA-seq detected expression of all three TET enzymes in Kiss1 neurons (data not shown). These observations suggest a potential role for Tet enzymes in regulating *Kiss1* gene expression and Kiss1 neuron function. We next performed hMeDIP-seq on DNA purified from the ARC of DIO mice and chow-fed controls, as previously described [25]. To minimize the impact of individual variation within each treatment group, we pooled equal amounts of DNA from three mice per group to generate biological replicates for hMeDIP-seq.

The 5hmC levels in the ARC were high at the gene body and correlated positively with gene expression (**Figure 6A**), consistent with previous reports [42]. 5hmC was depleted at the transcriptional start site (TSS) of actively transcribed genes, but was high at the TSS of repressed genes (**Figure 6A**). Differential analysis of 5hmC methylome in chow and DIO mice revealed widespread 5hmC changes in the ARC in these obese hyperglycemic male mice. We identified 94,614 differential 5hmC regions (DhMR, |FC|≥1.5 & p<0.001), spanning >60 million basepairs in total. Of these, 68,515 (72.4%) had 5hmC loss (HFD.down DhMRs) and 26,099 sites had increased 5hmC (HFD.up DhMRs) in DIO mice.

**Figure 6.**
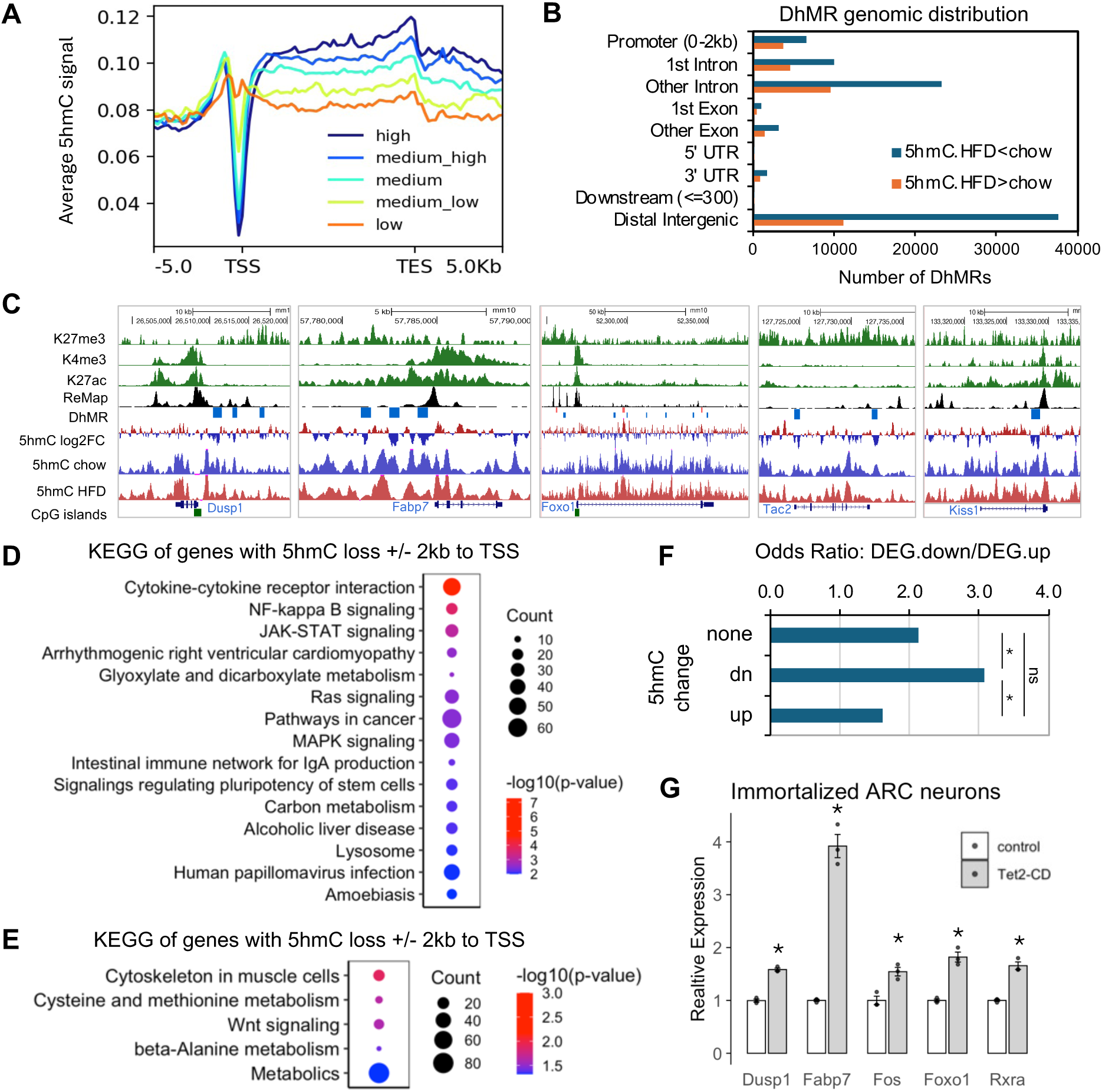
Reprogramming of the 5hmC methylome in the ARC of DIO mice. **A.** Metagene analysis of average 5hmC profiles at genes stratified by expression level. **B.** Genomic distribution of differential hydroxymethylation regions (DhMRs) with significant 5hmC increase or loss in DIO ARC. **C.** Representative 5hmC changes at key metabolic and neuronal genes. ReMap, density plot of transcription factor binding sites[78]. **D-E.** KEGG pathway enrichment of genes associated with DhMRs with significant 5hmC loss (**D**) or significant 5hmC increase **(E)**. The top 15 enriched pathways are shown for DhMRs with 5hmC loss in DIO ARC. **F.** The odds ratio of down- and up-regulated genes with no 5hmC changes (no DhMR association), with 5hmC loss (down), or with 5hmC increase (up) within TSS ± 2 kb. DhMR.down associated genes were more likely to be down-regulated compared to groups with no change or 5hmC gain. *, p<0.05, pair-wise proportion test following Chi square tests of 3 groups. **G**. Ectopic expression of the Tet2 catalytic domain (Tet2-CD) induces the upregulation of key metabolic and neuronal genes. Error bars, standard error of biological triplicates. *, p<0.01.

A significant portion of the DhMRs were located in the upstream promoters and in the first introns of genes (**Figure 6B**), close to the TSS (±2kb), accounting for 20.0% of HFD.down DhMRs and 26.2% of HFD.up DhMRs, respectively. Representative 5hmC changes are shown in **Figure 6C**. We identified 1911 genes with 5hmC loss close to the TSS (with DhMRs located within ±2kb to TSS), which are enriched in metabolism-related KEGG pathways, including glyoxylate and dicarboxylate metabolism and carbon metabolism, and inflammation-related pathways such as human papillomavirus infection and amoebiasis, as well as important signaling pathways such as NF-kappa B, JAK-STAT signaling, MAPK signaling, and cytokine-cytokine receptor interaction (**Figure 6D**). Nine hundred thirty-nine genes associated with increased 5hmC close to the TSS were enriched in KEGG amino acid metabolism, the WNT signaling pathway, and metabolic pathways (**Figure 6E**).

Examining the relationship of 5hmC changes (TSS±2kb) and differential gene expression, we found that DEGs with 5hmC loss close to a TSS were three times more likely to be down-regulated than up-regulated. The odds ratio was significantly higher than DEGs without 5hmC change or with 5hmC increase close to the TSS (**Figure 6F**, Chi square p<0.05). The preferential association of 5hmC loss with gene down-regulation is consistent with its known function in safeguarding low levels of DNA methylation at gene promoters and enhancers [43].

To examine the role of 5hmC in gene regulation directly, we transfected a plasmid encoding the catalytic domain of Tet2 (Tet2-CD) into KTaR cells, an immortalized cell line derived from ARC Kiss1 neurons [44], and examined effects on metabolic and neuronal gene expression using RT-qPCR. Without the N-terminus that promotes Tet2 protein degradation, the expression of the Tet2 catalytic domain robustly induced global increases in 5hmC in cells [45]. We observed significant up-regulation of *Dusp1*, *Fos*, *Fabp7*, *Foxo1*, and *Rxra* in Tet2-CD transduced cells (**Figure 6G**). *Dusp1*, dual specificity phosphatase 1, modulates neuronal excitability, synaptic plasticity, and has neuroprotective effects by limiting the overactivation of stress-related signaling, particularly in response to oxidative and metabolic stress [46]. *Fabp7* is the primary fatty acid transporter in the brain and modulates lipid signaling pathways, such as the NF-κB pathway, influencing inflammation and stress responses. *Fabp7* plays critical roles in both neurodevelopment and neuroplasticity, and its dysregulation is associated with Alzheimer’s disease [47]. c-fos, encoded by *Fos*, is considered a marker of neural activity [48]. *Dusp1*, *Fabp7* and *Fos* were significantly downregulated in the ARC of DIO mice and had significant 5hmC loss at the promoters (**Figure 6C**).

Although their expression was not altered in DIO ARC, we examined *Foxo1*, *Pparα*, and *Rxrα* given the dysregulation of PPAR signaling and lipid metabolism pathways revealed by our RNA-seq analysis and their association with DhMRs (**Figure 6C** and data not shown). *Foxo1* is a key transcription factor involved in mediating insulin signaling and plays a key role in regulating energy homeostasis in peripheral tissues [49] and in the hypothalamus [50]. PPARα and RXRα form a heterodimeric complex that is essential for regulating lipid metabolism [51]. In TET2-CD transduced cells, expression of the *Ppara* gene was not changed (not shown), whereas the *Rxra* and *Foxo1* gene expression were upregulated significantly. These genomic studies and functional validations strongly support a key function of 5hmC in regulating metabolic and neuronal gene expression in Kiss1 neurons.

## Discussion

Kisspeptin neurons are central in metabolic regulation of reproduction and play a role in regulating metabolic parameters, including food intake, glucose metabolism, respiratory rates, and thermoregulation [6, 52–54]. Recent studies have significantly expanded our understanding of the metabolic and epigenetic regulation of *Kiss1* gene repression or activation in the context of puberty development [8, 55]. In contrast, we know little about the mechanisms underlying the metabolic and epigenetic regulation of Kiss1 neuron function in adults. Our studies of well-characterized high-fat diet-induced obese diabetic mice demonstrated that Kiss1 neurons in the hypothalamic arcuate nucleus are key targets of the effects of metabolic disorders, sensitive to dietary and metabolic changes, and validated the role of suppressed Kiss1 neuron function as a key mechanism contributing to obesity/T2D-assocated male secondary hypogonadism. Epigenetic and transcriptomic investigation and functional validation have uncovered a key role of metabolism-induced 5hmC changes in regulating metabolic pathways in Kiss1 neurons in DIO mice. Taken together, our studies support a model in which Kiss1 neurons sense and respond to metabolic cues and consequently induce changes in the epigenome, metabolism, and function, thereby revealing a new mechanism underlying crosstalk between Kiss1 neurons and metabolism, in addition to regulation by upstream POMC/AgRP neuronal circuits that control energy homeostasis.

Our studies have demonstrated an impaired response of Kiss1 neurons to NK3R activation (i.e., senktide administration, in intact animals) and to testosterone depletion (delayed LH surge in GDX mice). Similarly, Villa et al. reported suppression of Kiss1 neuron activity in response to glutamatergic receptor activation and POMC neuron inputs [56], highlighting broadly reduced sensitivity to diverse stimuli. Although downregulation of *Kiss1* gene expression has been reported in HFD-fed animal models [37, 57], we did not detect significant changes in *Kiss1*, *Tac2*, or *Tacr3* gene expression, consistent with Villa et al. and others [56, 58]. Variable or absent Kiss1 changes have also been described in DIO female mice with disrupted estrous cycles and altered hormone profiles [59, 60]. Nevertheless, we observed significant 5hmC loss at the *Kiss1* gene locus in DIO mice (**Figure 6C**), suggesting metabolism-driven reprogramming of the local epigenetic landscape that may alter transcriptional responsiveness of the *Kiss1* gene in certain contexts. Importantly, our transcriptome analysis revealed pronounced alterations in metabolic and neuronal pathways, which likely underlie the changes in Kiss1 neuron function in the setting of obesity and type 2 diabetes mellitus, marked by diminished neuronal firing activity in response to inputs of diverse signaling pathways.

It is striking to observe the transcriptional downregulation of many genes encoding fatty acid transporters (including *Fabp1, Fabp7, Apoa1, Apoa5, Apoa2, Apoc1, Apoc3, Apoc4, Apof*, and others) and, concordantly, significant down-regulation of the PPAR signaling pathway and fatty acid metabolism pathway in the ARC of DIO mice. Neurons have limited capacity for fatty acid oxidation [61]. Hyperlipidemia and potential damage of the blood-brain barrier in obesity [62] are expected to result in increased uptake of fatty acids in the ARC [63]. The downregulation of fatty acid transporters and metabolism machinery in the ARC in DIO mice may contribute to ineffective clearance and accumulation of fatty acids in cells or in their microenvironment, leading to inflammation or cytotoxicity, which have a detrimental effect on neuronal function [64]. The association of 5hmC changes with key metabolic genes in the ARC in DIO mice, together with the ability of Tet to regulate expression of metabolic genes in hypothalamic cell lines, support a key role for metabolism-induced 5hmC changes in modulating Kiss1 neuron actvity, consistent with the emerging roles of 5hmC in PPARα [65] and PPARγ signaling [66, 67].

The role of 5hmC changes induced by alterations in metabolism in Kiss1 neurons is likely multifaceted, given the broad distribution of these changes in the genome. It is interesting to note the significant 5hmC alterations at key neuronal genes, including several immediate early genes. We show that early response genes such as *Jun* and *Fos* are not only down-regulated in the ARC of DIO mice, they are also direct targets of 5hmC in Kiss1 neuronal cell lines. Under normal conditions, IEGs are activated rapidly in response to neuronal activation, and subsequently rapidly inactivated, making them a useful indicator of recent activation. It has been reported that DIO mice have fewer c-fos positive Kiss1 neurons [56]. Sustained upregulation of IEGs was associated with the long-term increase in 5hmC at these genes in mouse models of cognitive behavioral abnormalities [68], supporting the functional importance of 5hmC in IEG downregulation and Kiss1 neuron dysfunction in DIO mice *in vivo*.

The loss of 5hmC at the majority of DhMRs (>70%) is consistent with the control of 5hmC levels via the glucose-AMPK-TET2 signaling pathway [11]. Hyperglycemia is a key aggravating factor that is independently associated with male hypogonadism. Intracranial glucose levels correlate with blood glucose fluctuations, albeit with a dampened response in patients with type 2 diabetes [69]. In longitudinal studies, type 2 diabetes was associated with lower total testosterone levels, after adjusting for age and body mass index (BMI) [70, 71]. Obesity is frequently associated with elevated blood glucose levels in patients and in animal models. In our DIO cohorts, we observed a significant increase in fasting plasma glucose levels as soon as 4 weeks after initiation of a high fat diet, and levels continue to increase in association with additional weight gain. We speculate that hyperglycemia may induce 5hmC loss via the AMPK-TET2 signaling pathway, and consequently causes dysregulation of the PPAR signaling pathway and fatty acid metabolism in Kiss1 neurons, consistent with the reported function of TET2 in fatty acid metabolism, adipocyte development, and obesity [10, 65, 66]. Tet2⁻/⁻ mice exhibited an age-associated progressive decline in fecundity [72]. Though the reduced litter size was attributed to defects in meiosis in the oocytes of aged animals, reduced ovarian weights were observed in young and middle-aged (<9 month old) Tet2^-/-^ mice before the onset of oocyte deterioration [72], suggesting reproductive defects in addition to accelerated oocyte aging, which warrants further investigation.

It is important to note that the mechanisms of metabolic regulation of TET activity are complex [9, 73]. Sirt1 (Sirtuin 1), an NAD^+^-dependent deacetylase suppressed by overnutrition, is important for pre-pubertal repression of *Kiss1* gene expression and impacts the timing of puberty onset in female mice in response to energy state [7]. Sirt1 also regulates Tet2 protein stability by preventing post-translational acetylation of the C-terminal domain of Tet2, which targets Tet2 to protein degradation pathways [74]. In addition to the glucose-AMPK-TET2 signaling pathway, it is reported that high glucose inhibits TET3 activity by increasing TET3 *O*-GlcNAcylation, which drives TET3 nuclear exclusion *ex vivo* in cultured cells [75]. TET1/2/3 play complementary roles in neuron biology [76] and metabolism [66, 77]. Thus, multiple mechanisms may contribute to 5hmC loss in the ARC of DOI mice. Future studies need to validate the key functions of Tet2 and other Tet enzymes in regulating Kiss1 neuron metabolism and function in obesity and type 2 diabetes, and their interplay with AMPK and PPAR signaling.

## Conclusion

Taken together, we have identified suppressed Kiss1 neuron function as a key mechanism underlying obesity-associated male secondary hypogonadism in a high-fat diet–induced obese diabetic mouse model. Mechanistically, we have demonstrated that Kiss1 neurons in the arcuate hypothalamus are highly sensitive to dietary and metabolic cues, and that metabolic disorder–induced 5hmC changes play an important role in altered gene expression and the dysregulation of Kiss1 neuron metabolism and function. These findings reveal a novel mechanism linking metabolic disturbances to reproductive dysfunction through direct effects on Kiss1 neurons, independent of upstream POMC and AgRP inputs.

## Data availability

The datasets supporting the conclusions of this article are available in the GEO repository (nos. **pending**).

## Abbreviations

5hmC: 5-hydroxymethylcytosine
5mC: 5-methylcytosine
AgRP: agouti-related peptide
AMPK: adenosine 5′-monophosphate (AMP)-activated protein kinase
Apo: apolipoprotein
ARC: arcuate nucleus hypothalamus
DEG: differential express genes
DhMR: differential hydroxymethylation region
DUSP1: dual specificity phosphatase 1
Fabp: Fabp
Foxo1: forkhead box O1
FSH: follicle-stimulating hormone
GDX: gonadectomy
GLP-1: glucagon-like peptide-1
GOBP: gene ontology biological processes
hMeDIP: hydroxylmethylation DNA immunoprecipitation seqeuencing
HPT: hypothalamic–pituitary–testicular
IEG: immediate early gene
KEGG: Kyoto Encyclopedia of Genes and Genomes
Kiss1: kisspeptin
KNDy: kisspeptin, neurokinin B, dynorphin co-expressing neurons
LH: luteinizing hormone
NK3R: neurokinin 3 receptor
NKB: neurokinin B
POMC: pro-opiomelanocortin
PPAR: peroxisome proliferator–activated receptor
Rxra: retinoid X receptor alpha
SEM: standard error of measurement
Sirt1: Sirtuin 1
T2D: type 2 diabetes
Tac2: tachykinin 2
Tacr3: tachykinin receptor 3
Tet2-CD: TET2 catalytic domain
TET2: tet methylcytosine dioxygenase 2
TSS: transcription start site

## Acknowledgments

The authors would like to thank Han Kyeol Kim and Melissa Magnuson for their technical assistance. The KaTR cell line was a generous gift from Dr. Patrick Chappell.

## Author information

### Author notes

Rui Fang and Ursula B. Kaiser are co-communicating and co-senior authors and contributed equally to this work.

## Contributions

Conceptualization: RF and UBK; Methodology: RF, RSC and UBK; Investigation: RF; Data Analysis: RF; Writing - Review & Editing: RF and UBK; Supervision: RSC and UBK; Interpretation of Results: RF, RSC and UBK.

## Ethics declarations

### Ethics approval and consent to participate

Not applicable.

### Consent for publication

Not applicable.

