## Supplemental Table and Figures for "High-Fat Diet Induces Epigenetic and Metabolic Changes in Kisspeptin Neurons in Association with Obesity and Male Secondary Hypogonadism"

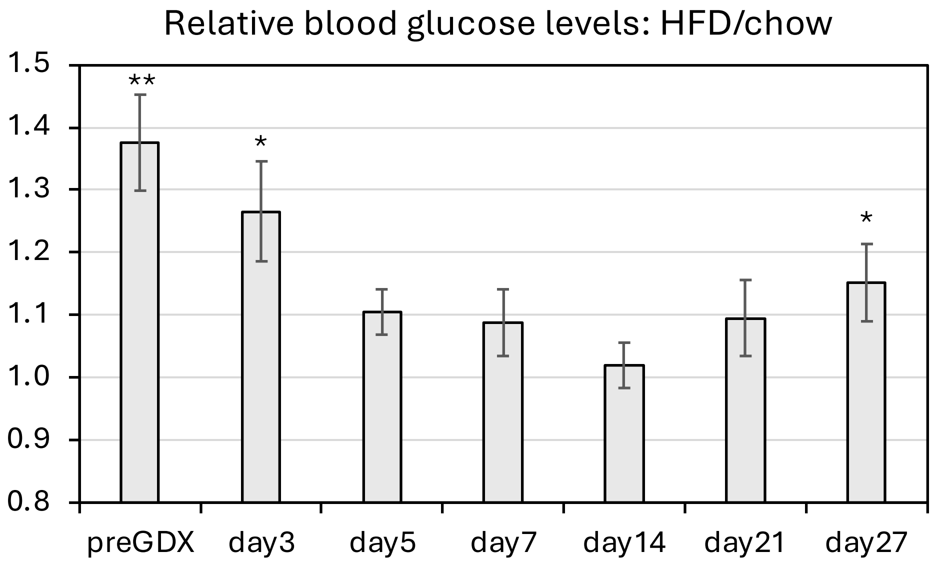


**Fig S1. Blood glucose in chow and HFD mice post GDX surgery.**
Fold change of blood glucose in HFD mice relative to the mean of chow group were shown. HFD, n=5; chow, n=6. Error bars, SEM. *, $p<0.05$; **, $p<0.01$; Student's *t*-test.

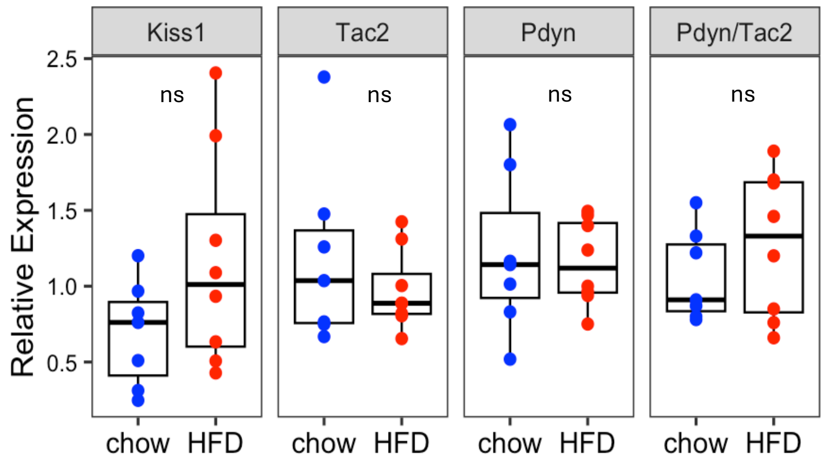


**Fig S2. Gene expression of *Kiss1*, *Tac2* and *Pdyn* were not changes in the ARC of HFD-induced obese mice.**
qRT-PCR was performed on ARC tissue from intact male mice fed a high-fat diet (HFD) for 16 weeks and age-matched controls fed standard chow. Expression was normalized to GAPDH. Chow, n=7; HFD, n=8. In boxplot, line within box denotes the median. ns, not significant by Student's *t*-test.

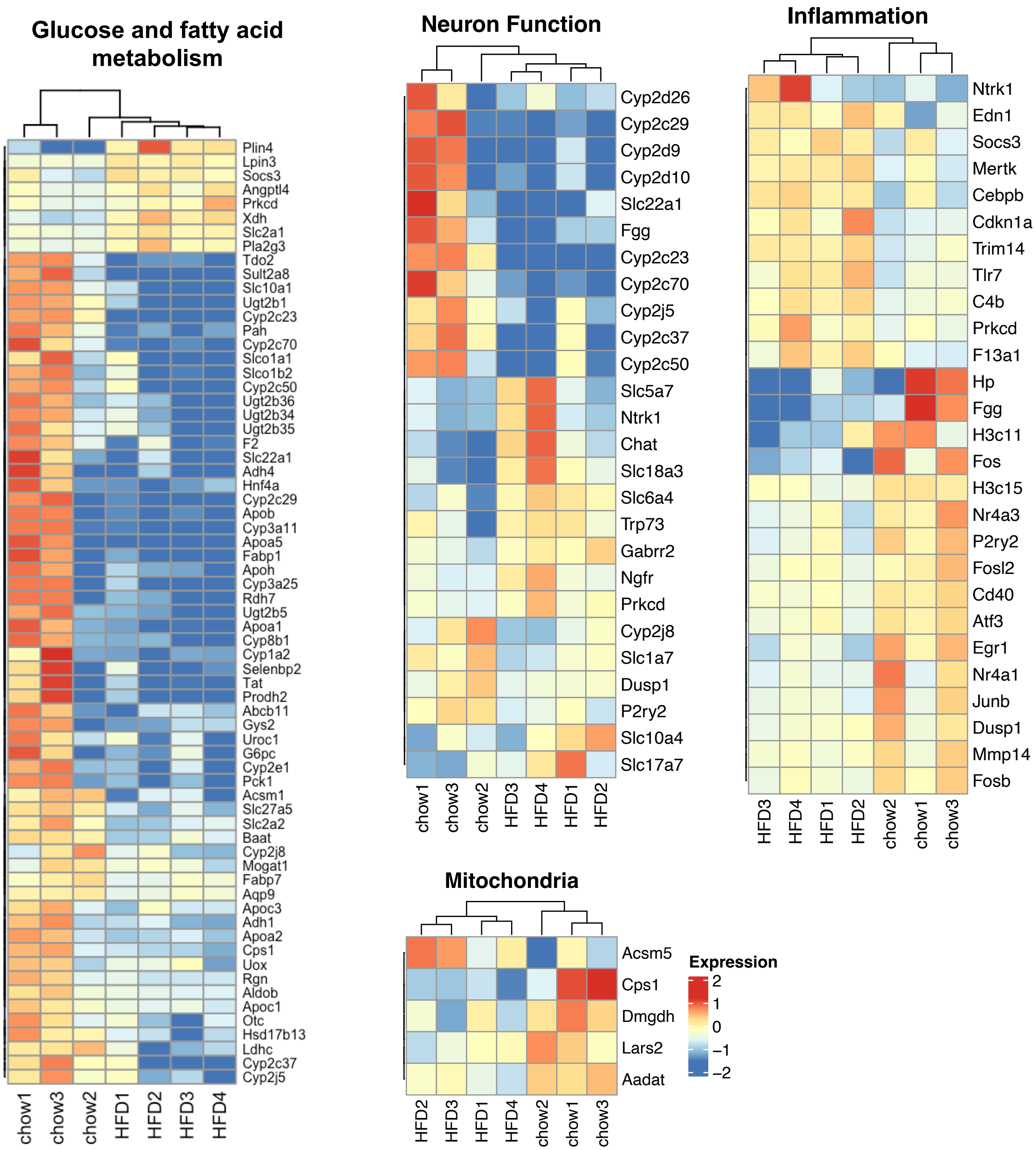


**Fig S3. Heatmap of differential gene expression in the ARC of HFD-induced obese mice compared to mice fed standard chow.**

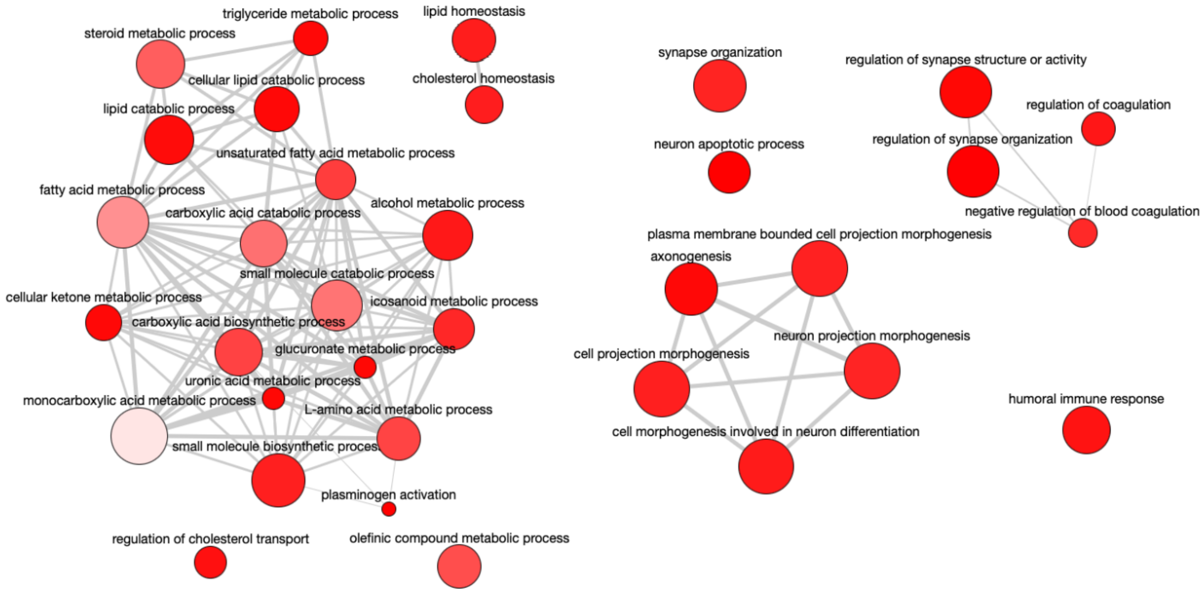


**Fig S4. GOBP terms identified by GSEA in ARC transcriptomes of HFD- and chow-fed mice.**

Reduce representation of significantly enriched GOBP terms (FDR<0.05) were plotted using REVIGO[1]. Bubble color indicates p-value; bubble size indicates the frequency of the GO term. Highly similar GO terms are linked by edges in the graph, where the line width indicates the degree of similarity.

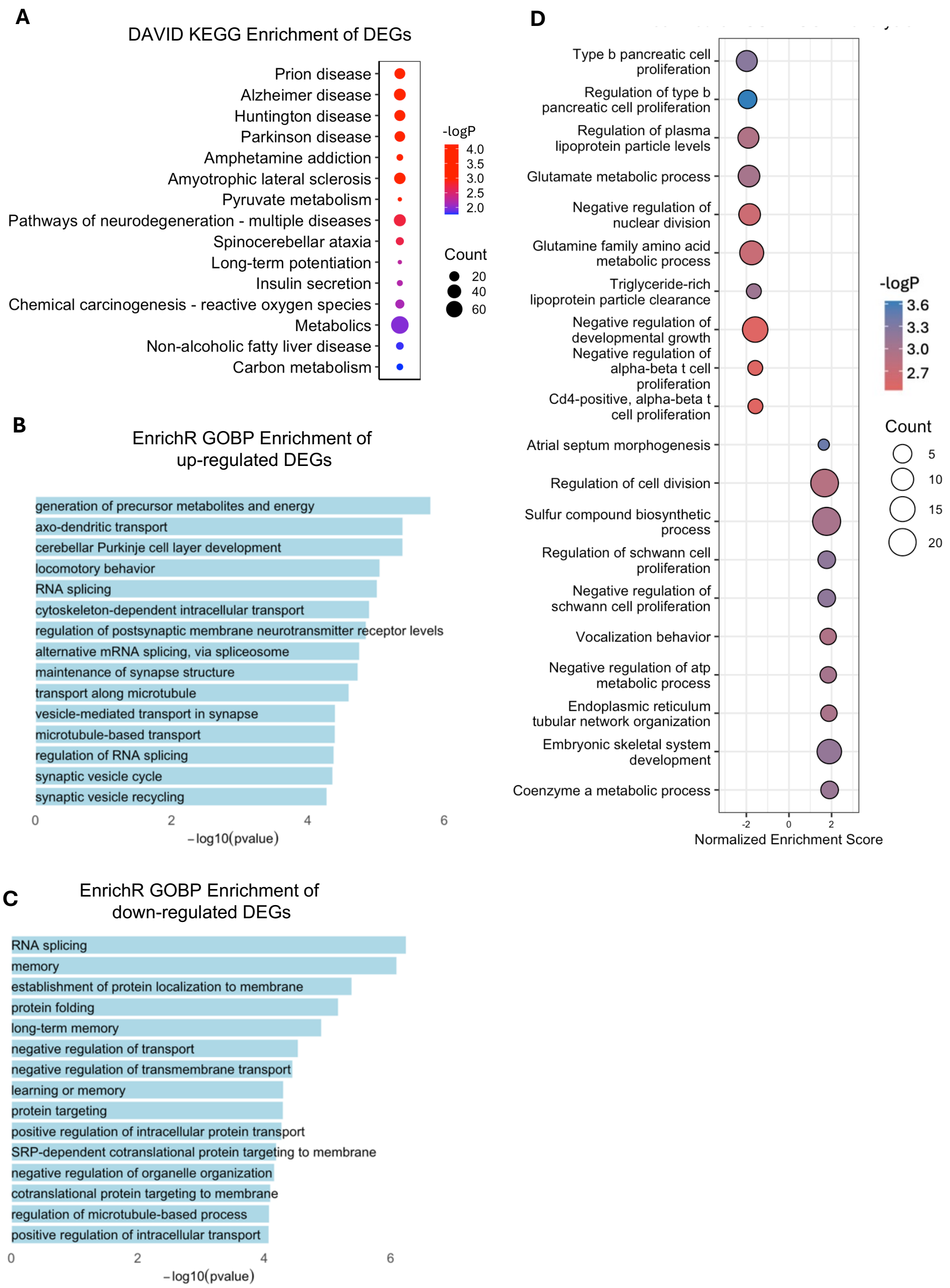


**Fig S5. KEGG and GOBP pathways enrichment among differentially expressed genes in Kiss1⁺ neurons from mice fed on LFD or HFD.**

**A.** KEGG gene pathway enrichment of DEGs identified using a relax cutoff of |FC|≥1.5 and p<0.1 in single-cell RNAseq dataset (Campbell, et al [40]).

**B, C.** GOBP enrichment of up-(**B**) and down-(**C**) regulated DEGs using EnrichR[2].

**D.** Top 10 most up- and down-regulated GOBP pathways by GSEA.

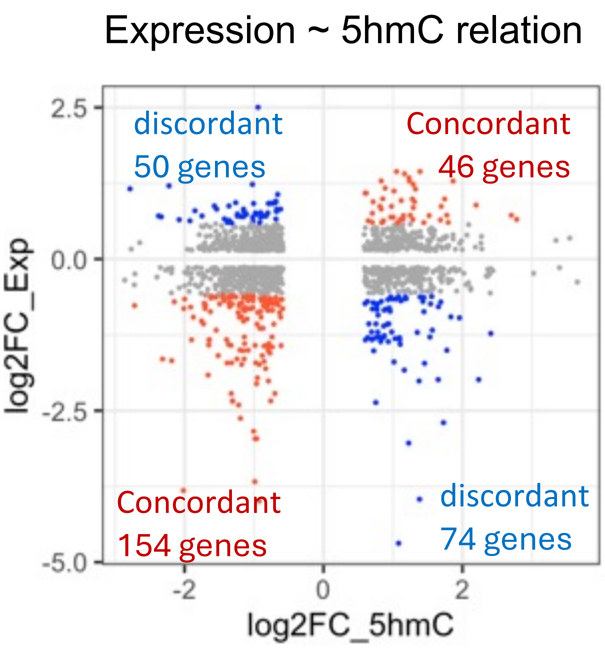


**Fig S6. The correlation of 5hmC changes and gene expression changes in the ARC of mice fed chow or HFD.**

There are 3469 DhMRs (2761 5hmC HFD>chow; 1280 5hmC HFD<chow; |FC|≥1.5 & $p<0.001$ & FDR<0.05) associated with 2761 genes (±2kb to TSS). Of these, 1461 genes with ≥1.1 fold changes in gene expression were shown in the plot. Genes with less than 1.1 fold change in expression or with very low expression (RPKM<0.05) were excluded. Grey, less than 1.5 and greater than 1.1 fold change in gene expression. Red or blue, greater than 1.5 fold change in gene expression. Red, concordant changes in 5hmC and gene expression (both increase or both increase). Blue, discordant changes in 5hmC and expression (increase in 5hmC and decrease in gene expression or vice versa). Each dot represents one gene. The number of DEGs (≥1.5 fold changes) in each group was shown.

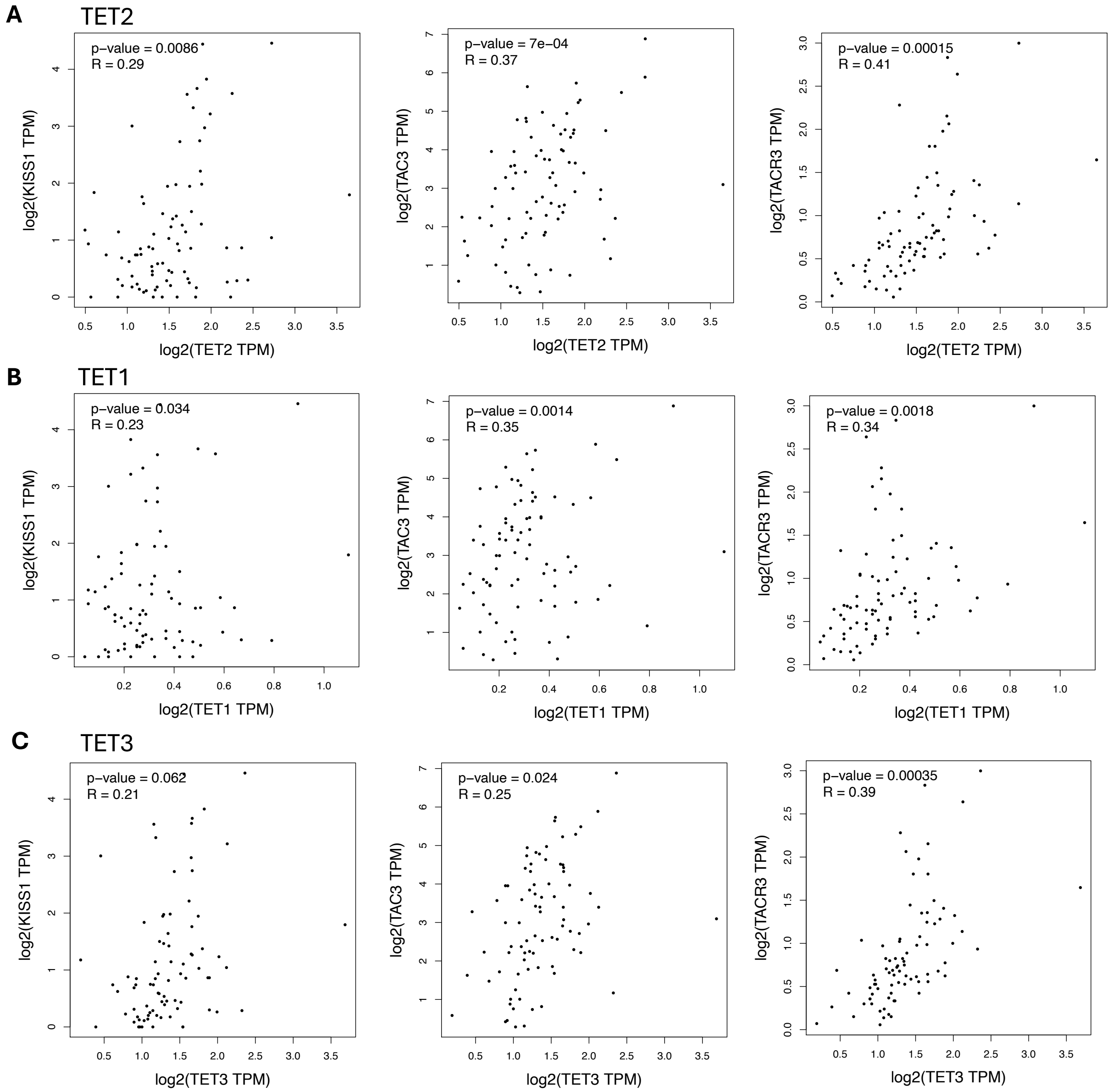


**Fig S7. The Correlation of *TET1*, *TET2*, and *TET3* gene expression with *KISS1*, *TAC2* and *TACR3* in human hypothalamus**

Correlation analyses were performed on healthy hypothalamic samples from the GTEx database using using GEPIA[3]. Pearson's correlation coefficient (R) and p value were shown.

**Supplementary Table 1. Sequence of RT-qPCR primers**

| **Gene** | **Forward Primer Sequence** | **Reverse Primer Sequence** |
|---|---|---|
| Kiss1 | 5’-TGATCTCAATGGCTTCTTGG-3’ | 5’-CCCAGGCATTAACGAGTTC-3’ |
| Tac2 | 5’-TGGAAGGATTGCTGAAAGTG-3’ | 5’-CCATAAGTCCCACAAAGAAG-3’ |
| Pdyn | 5’-CTCGTGATGCCCTCTAATG-3’ | 5’-CACTCCAGGGAGCAAATC-3’ |
| Gapdh | 5’-TGGAGTCTACTGGTGTCTTC-3’ | 5’-ACACCCATCACAAACATGG-3’ |
| Dusp1 | 5’-CTGAGTACTAGTGTGCCTGAC-3’ | 5’-GTACTGGTAGTGACCCTCAAAG-3’ |
| Fabp7 | 5’-GGACACAATGCACATTCAAG-3’ | 5’-CAACCGAACCACAGACTTAC-3’ |
| Fos | 5’-GCCTTTCCTACTACCATTCC-3’ | 5’-AAAGTTGGCACTAGAGACG-3’ |
| Foxo1 | 5’-ACGAGTGGATGGTGAAGAG-3’ | 5’-GCTGTGAAGGGACAGATTG-3’ |
| Rxra | 5’-CTGTTCAACCCTGACTCTAAG-3’ | 5’-CGCTTCTAGTGACGCATAC-3’ |

1 Supek, F., Bosnjak, M., Skunca, N. & Smuc, T. REVIGO summarizes and visualizes long lists of gene ontology terms. *PLoS One* **6**, e21800 (2011). https://doi.org/10.1371/journal.pone.0021800

2 Chen, E. Y. *et al.* Enrichr: interactive and collaborative HTML5 gene list enrichment analysis tool. *BMC Bioinformatics* **14**, 128 (2013). https://doi.org/10.1186/1471-2105-14-128

3 Tang, Z. *et al.* GEPIA: a web server for cancer and normal gene expression profiling and interactive analyses. *Nucleic Acids Res* **45**, W98-W102 (2017). https://doi.org/10.1093/nar/gkx247
